# Microscope-Cockpit: Python-based bespoke microscopy for bio-medical science

**DOI:** 10.1101/2021.01.18.427178

**Authors:** Mick A Phillips, David Miguel Susano Pinto, Nicholas Hall, Julio Mateos-Langerak, Richard M Parton, Josh Titlow, Danail V Stoychev, Thomas Parks, Tiago Susano Pinto, John W Sedat, Martin J Booth, Ilan Davis, Ian M Dobbie

## Abstract

We have developed “Microscope-Cockpit” (Cockpit), a highly adaptable open source user-friendly Python-based GUI environment for precision control of both simple and elaborate bespoke microscope systems. The user environment allows next-generation near-instantaneous navigation of the entire slide landscape for efficient selection of specimens of interest and automated acquisition without the use of eyepieces. Cockpit uses “Python-Microscope” (Microscope) for high-performance coordinated control of a wide range of hardware devices using open source software. Microscope also controls complex hardware devices such as deformable mirrors for aberration correction and spatial light modulators for structured illumination via abstracted device models. We demonstrate the advantages of the Cockpit platform using several bespoke microscopes, including a simple widefield system and a complex system with adaptive optics and structured illumination. A key strength of Cockpit is its use of Python, which means that any microscope built with Cockpit is ready for future customisation by simply adding new libraries, for example machine learning algorithms to enable automated microscopy decision making while imaging.

**Highlights:**

- User-friendly setup and use for simple to complex bespoke microscopes.
- Facilitates collaborations between biomedical scientists and microscope technologists.
- Touchscreen for near-instantaneous navigation of specimen landscape.
- Uses Python-Microscope, for abstracted open source hardware device control.
- Well-suited for user training of AI-algorithms for automated microscopy.

## Introduction

### Why we need Cockpit

Biomedical research has benefited from significant engineering advances that have increased the speed and sensitivity of many types of scientific instrumentation. Microscope technology, in particular, is evolving rapidly through advances in optical techniques and image processing, enabling major scientific discovery. However, the rapid adoption and application of advanced microscopies by biomedical scientists is constrained, especially when it relies on commercialisation to make the necessary technology accessible to the end-users. It can often take several years to develop innovations from research instruments into user-friendly, off-the-shelf systems[1, 2, 3, 4]. We report a collaboration between the University of Oxford and UCSF that grew out of the OMX microscope project, dating from the early 1990’s, which is extensively documented in the Supplement of [1].

To accelerate technology adoption, many labs are building custom microscopes based on newly developed imaging methods, such as light sheet[5], lattice light sheet [6], 3DSIM [1, 7], STED [8], and MINFLUX [9], or to exploit new analysis techniques, such as STORM analysis[10, 11], 3B[12], and SIMFLUX [13]. These instruments require flexible, computerised control of a wide range of individual components (lasers, mirror actuators, filters, objectives, detectors, etc.) allowing more rapid deployment of novel techniques, and often providing faster and more sensitive operation than commercial systems[6, 14, 15]. However, integration of these components into a single user-friendly platform that is focused on users’ scientific application, rather than dictated by the control infrastructure, can be a major challenge. There have been three general solutions adopted by the community for software control of bespoke microscope hardware.

1. Using individual manufacturer-provided packages for each piece of hardware. This can provide a straightforward solution for simpler systems, but fundamental incompatibilities between each vendor’s control models can prevent integration of multiple components.
2. Using LabVIEW, a commercial visual programming tool from a leading hardware manufacturer, National Instruments.
3. Use of a dedicated microscope control package, of which several are available, with the most widely used being Micro-manager (μManager): an open source Java and C++ based software platform for controlling a range of microscope hardware such as stands, cameras, and stages. A less widely-adopted solution is a bespoke platform written in Python to control the OMX microscope [1, 16, 17], and this forms the starting point of the Microscope-Cockpit project described in this manuscript.

An important feature for custom microscopes is the freedom to design the system around a specific set of experiments, as defined by the user. With a specific application in mind, an instrument can be optimised to make the best possible compromises for that application. For example, imaging deep into biological tissue can involve significant optical aberrations leading to degraded image quality. This can be compensated for by the application of adaptive optics techniques, at a cost of increased complexity and marginally decreased light efficiency. However, it is desirable that this capability is easily enabled and exploited by microscope users, typically biologists with limited experience of the underlying engineering, and be reproducible over months or years of use. Many such custom microscope designs have been published [18, 19, 20, 21, 22, 23]. In all of these cases, control of the many hardware components in a complex and timed manner is challenging, particularly for time sensitive applications involving living specimens. Correction settings must be measured and applied in coordination while simultaneously minimising additional exposure of the sample.

Finally, the democratisation of AI-based image analysis approaches means that a wealth of tools are potentially available to users. However, current systems, with some specific exceptions, are not well suited for the integration of bespoke novel AI-approaches with the imaging process. This limitation can be significantly reduced by an appropriately flexible package for control of bespoke systems.

### Philosophy and implementation of Cockpit

We set out to create a package that is suitable for biologists to carry out a range of experiments in a time efficient manner, at scale. At the same time, the software has to control a wide range of electronic and optical hardware devices and microscope types. Many bespoke microscope systems that lack eyepieces, such as systems built by physical scientists in collaboration with biologists, making sample navigation particularly challenging. Improving navigation on this type of system is a major feature of Cockpit.

There are several existing software control approaches including manufacturer-supplied packages, custom control software, in LabVIEW or a more traditional programming language, or the open source program μManager, as discussed earlier. We feel that Cockpit has several distinct advantages over these other approaches. We compare the relative merits of Cockpit and a range of alternatives in the discussion.

Cockpit is an open-source software package that makes it easy to control multiple devices through a single Python-based user interface (Figure 1). Cockpit supports multiple platforms, and the similarity of its interface across platforms is shown by the main window in Linux, macOS, and Windows 10 Figure 2. It provides an innovative mosaic tool, allowing large areas of a microscope specimen to be imaged and then navigated in real time. This is similar to Micro-Magellan [24], but with the additional feature of keeping even thousands of images in GPU memory for instant access at various levels of detail. It utilises Python-Microscope (https://www.python-microscope.org) to communicate with a wide range of microscope hardware and is able to exploit a master timing control device to provide hardware triggering for digital devices and synchronised analogue voltages. This enables highly reproducible timing over a range of time scales even in extremely complex experiments with multiple devices.

**Figure 1:**
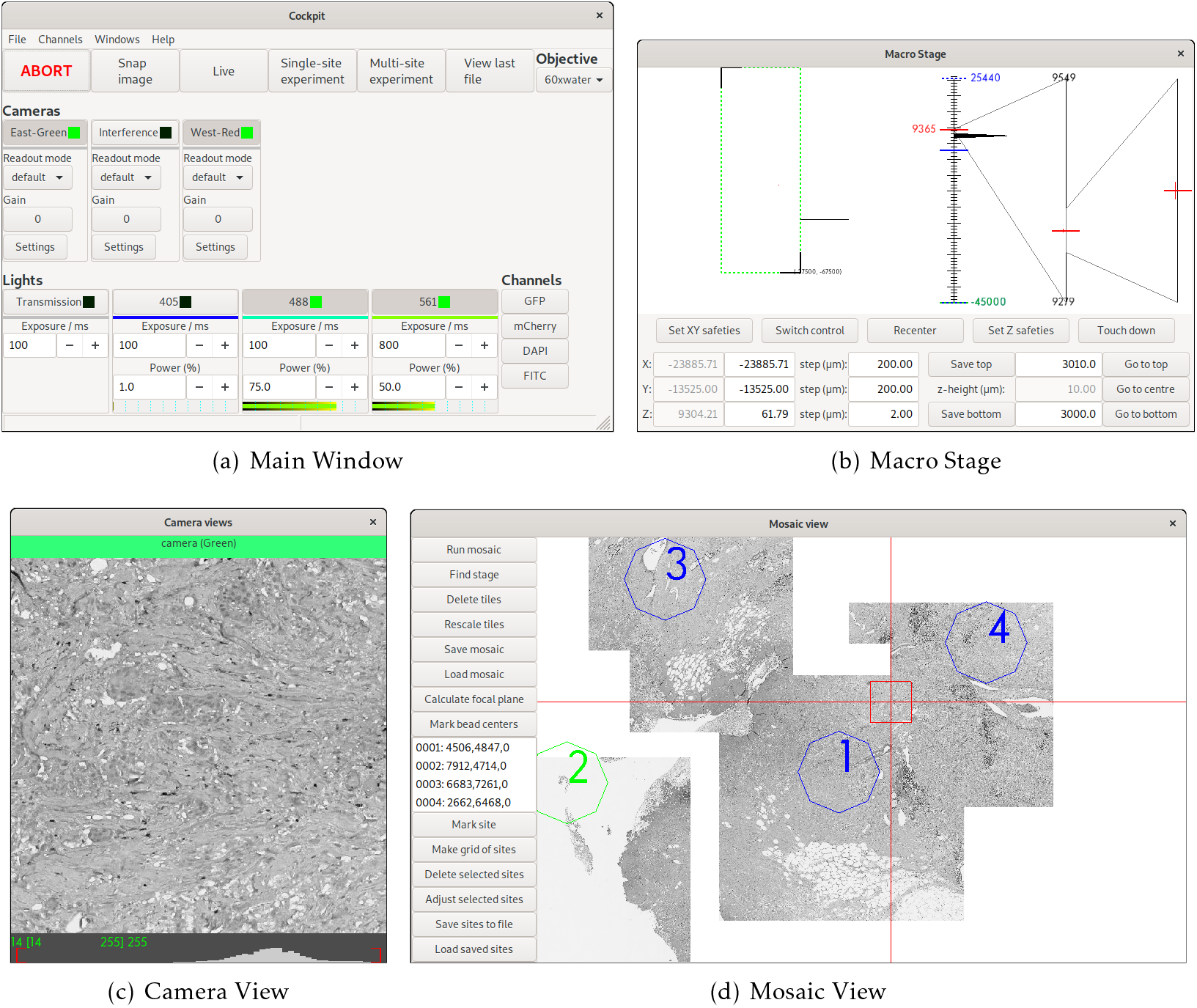
The main Cockpit GUI components: (a) The main window provides quick access to a number of functions and shows the status of all the devices, as well as the channels buttons to load predefined configurations. (b) The macro stage window provides an overview of the position of all stages, including nested stages and Z position. (c) The camera view shows the last image taken from each active camera and their histograms. (d) The mosaic view displays all mosaic images taken, saved locations, and dramatically eases navigation.

**Figure 2:**
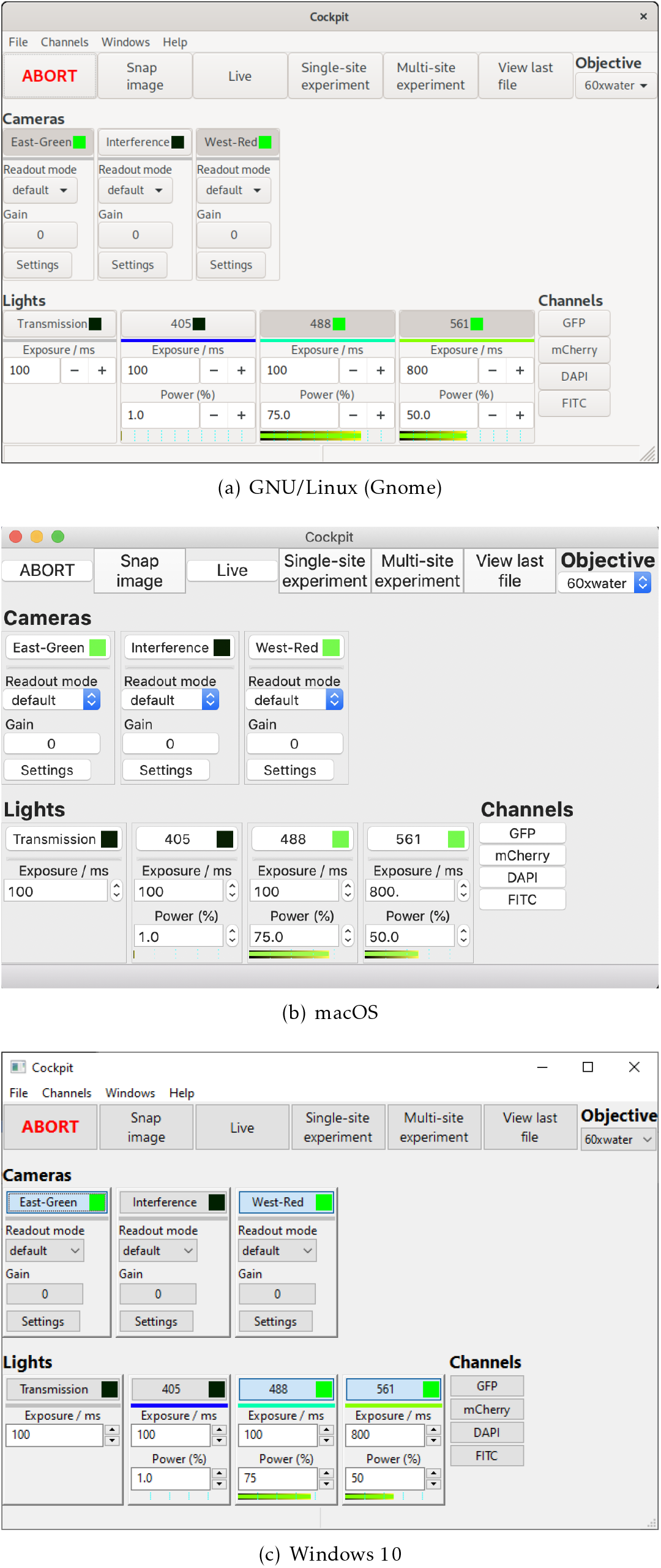
The Cockpit main window under different operating systems.

The simple interface design (Figure 1) enables users to focus on collecting the data they require without having to devise complex control infrastructure for systems of any level of complexity. It also enables easy navigation of even large sample regions. This can be especially important on bespoke microscopes which often lack eyepieces. For these types of system, Cockpit includes a touchscreen-optimised display window, providing large buttons and simple control over most functions (Figure 3).

**Figure 3:**
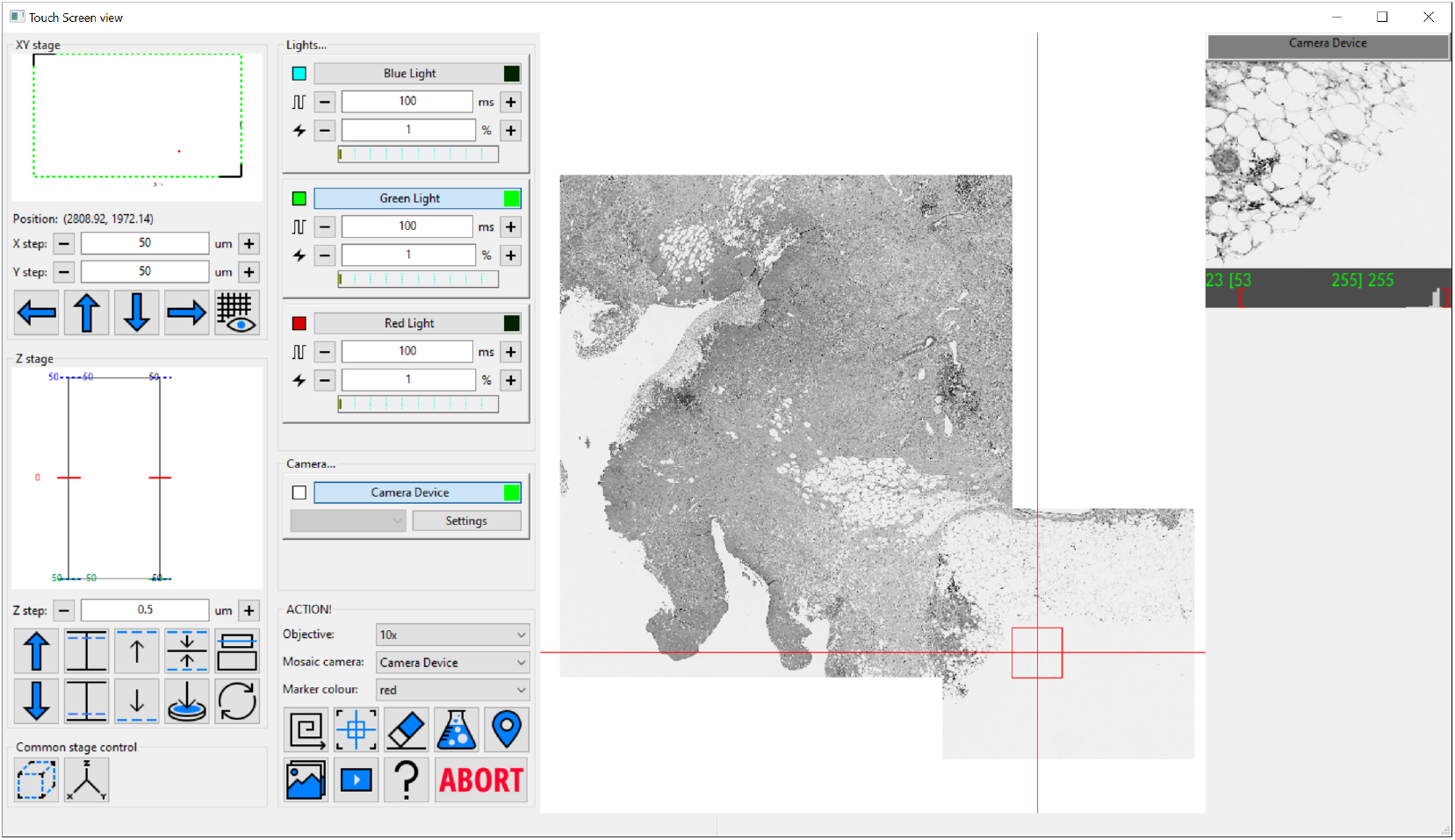
The Cockpit touchscreen window with the main GUI components in one window and large touchscreen friendly buttons. The left panel houses stage position indicators and controls, next panel over has devices like cameras and lights along with a set of large buttons at the bottom to run mosaics, centre the field of view on the current stage position, erase mosaic images, open the experiment dialog, drop a marker at the current position, display live video, run an experiment, a help button, and finally abort which stops an experiment or mosaic. The majority of the window is the mosaic display panel which can zoom from an overview up to the level of single pixels. Finally on the right is the last snapped image from each enabled camera, only one in this case.

The principal design goals for the Cockpit interface were to:

- Provide an easy-to-use, simple “Cockpit” control program for a wide range of microscopes.
- Hide from the user the complexity required to control an advanced microscope.
- Minimise the effort required to change hardware or integrate new hardware.
- Enable high-precision hardware timing signals to optimise experimental control, facilitate complex systems and maximise speed.

Cockpit provides an integrated microscope control package that enables all these features while being user friendly and portable across many different microscopes and common operating systems.

The first version of the Cockpit software was developed in UCSF to drive the OMX microscope [1, 17, 16], which was later commercialised. This code was re-written at UCSF and later released as open source under a BSD license. From this concept we have extensively developed the implementation of Cockpit to realise its full potential.

## Implementation

Cockpit is implemented in the Python programming language. The code consists of four main components: devices, handlers, interfaces, and the GUI. Devices represent the individual physical devices, handlers represent control components of the devices, handlers from single or multiple devices are aggregated into interfaces, and the GUI component allows user interaction with the different interfaces.

### Devices and Handlers

The hardware control side of Cockpit has two parts: devices and handlers. A Cockpit device maps to a single physical device while handlers abstract the control components of the device to be used. For example, an XY stage device provides two handlers, one per axis. A laser device also provides two handlers, a light source handler and a light power handler: the light source handler controls the light source state — on or off — while the light power handler controls its intensity.

Most of the Cockpit code does not interact with Cockpit devices directly. Rather, when the Cockpit program starts, it constructs a “depot” object whose role is to obtain and keep track of the different device handlers. This architecture enables rapid changes in the choices of hardware devices, so they can be very easily added or removed from a complex system as it is being developed. This approach prevents potential conflicts or confusions such as addressing devices that do not exist or adding devices that lack drivers.

Within this architecture, any Python class can be used as a device; it is only required to provide one or more Cockpit handlers. Built-in Cockpit devices include adaptors for the Python-Microscope package, thus supporting all of its available devices. Cockpit also supports a series of different stages, a spatial light modulator (SLM) from Meadowlark Optics, a polarisation rotator, and the SRS SR470 shutter.

Cockpit itself does not require a direct connection to the hardware. Indeed, most Cockpit devices are implemented as remote objects with the help of Pyro, a Python package for remote procedure calls. The use of Pyro enables the distribution of devices over the network to different computers, even running different operating systems.

### Interfaces and GUI

Cockpit defines interfaces that are high-level abstractions for the system, with each controlling one or more handlers. For example, the stageMover interface is a unified translation manager that coordinates all translation stages, hiding the complexity of handling different stages for different axes and nested stages on the same axis. For instance, a system may include an XY stage, a coarse Z stage, and a high precision fast piezo Z stage. The GUI component provides different views onto the multiple interfaces and handlers, each exposing different levels of complexity.

Cockpit has a multi-window GUI, with each window serving a different purpose (Figure 1). This provides the flexibility to distribute the windows over multiple monitors in whatever form best fits the system being used. The main window provides an at-a-glance view of the state of the various devices; the mosaic window enables navigation of the sample relative to an overview formed of tiled images; the macro stage window displays the position of the different stages; and the camera view the current images from all active cameras. In addition, Cockpit also has a simpler single-window touchscreen interface (Figure 3) which duplicates a subset of the functionality provided on the other windows. This window is optimised for touchscreen interaction with large buttons for the most useful subset of functionality.

Cockpit uses wxPython, a Python wrapper to wxWidgets, to provide a user interface that has a native look and feel on the different platforms (Figure 2). Within wxPython, we make extensive use of OpenGL to enable high performance display and interaction with large images or image mosaics up to gigapixel sizes.

### Data Acquisition Experiments

In Cockpit, the acquisition of an image series, such as a Z-stack or a time lapse sequence, is defined by an experiment. Cockpit experiments are based around the concept of an action table which must be run with hardware triggers. The experiment settings are converted into a list of device actions, which is parsed into a list of pre-computed analogue values and digital levels for each time point on a timing device. The action table is uploaded to the timing device and then started, running the experiment utilising hardware triggers and analogue voltages, where appropriate, to control all active devices. This approach, adapted from Carlton et al. [1], is able to produce both digital triggers and analogue signals with precise timing.

Effective execution of the experiment requires accurate timing. In order to ensure that experiment timing is independent of the host computer’s performance or the Python implementation, the timing functionality is performed using dedicated hardware. This enables time-critical operation of the microscope imaging tasks. The timing system has been implemented on several different hardware platforms, a digital signal processing board (Innovative Integration M67-160 with an A4D4 daughter board), an NI FPGA board (National Instruments cRIO-9068), and a Red Pitaya single board computer (STEMlab 125-14). In this way, we have created a universal and adaptable microscope platform with outstanding timing precision and accuracy.

### System Requirements

Cockpit requires Python 3.5 or later, and the Python packages PyOpenGL, Pyro4, freetypepy, matplotlib [25], microscope, NumPy [26], pySerial, SciPy [27], and wxPython. These are all free and open-source software that are available for all widely-used operating systems. Hence, Cockpit can be used across GNU/Linux, macOS, and Windows.

The Cockpit interface incorporates some assumptions as to what is required for a minimal microscope. It requires at least one camera, a computer controlled X, Y and Z translation motorised axes, and a light source that can either be hardware triggered (directly or via a shutter) or left on during an experiment. In addition, experiments require a device that can be programmed as source of hardware triggers, supplying digital signals and, optionally, analogue voltages.

With no configuration to specific hardware, Cockpit will automatically start in simulation mode, providing simulated cameras, light sources, and stages. This allows testing and demonstration of the software without any hardware or configuration. This can be further extended by utilising the test devices within the associated Python-Microscope package. In this mode, with some configuration, the software can link simulated XYZ stage, camera, and filter-wheel to return a channel defined by the filter-wheel and a subregion defined by the stage position from a large multi-channel image. The simulation is further improved by blurring the image based upon the Z position of the stage. This mode was used for many of the figures provided in this manuscript (Figures 1 & 3).

## Use Cases

Cockpit is optimised to provide an effective user experience running a fully automated microscope on systems with no eyepieces. A significant component of this are powerful mapping and overview tools that provide a guide-map to the specimen landscape, which is interactive and highly responsive. The system can collect arbitrary mosaic images, which are acquired in an expanding spiral pattern. Acquired mosaic images are immediately uploaded to the graphics card as a texture map. This allows smooth and rapid display of these images from low zoom overview all the way to single pixel views. The graphics card can then render these images interactively in real time. The typical texture memory on modern graphics cards mean that the system can navigate in real time around mosaics made from hundreds or thousands of individual images, with resulting mosaics up to gigapixels in size. Starting a new mosaic does not remove previous images, so multiple, possibly overlapping, mosaic areas can be collected. Keeping previous mosaic images means it is easy to navigate multiple regions of the sample without having to reacquire overview images. This greatly eases the navigation of complex samples.

There are two obvious use cases for this functionality. In the first case, one could utilise a low magnification lens to map a large section, or even the whole of the sample area, say a coverslip as shown in Figure 4(a). This produces a ~250 megapixel image, via which the user can visit any site of choice and re-image points in fluorescence mode with higher magnification objectives (Figure 4(b)). The new detailed images are then “painted” onto the guide-map in real-time. As the second case, in samples such as tissue culture cells, where many similar regions are available, smaller regions can be imaged at higher resolution to select interesting targets for further imaging. The preservation of previous mosaics when a new one is initiated is a critical component of this process, as it allows easy navigation of multiple sample regions. It is then practical to return to the regions of most interest without requiring reimaging, which would waste time and incur unnecessary light exposure to the sample.

**Figure 4:**
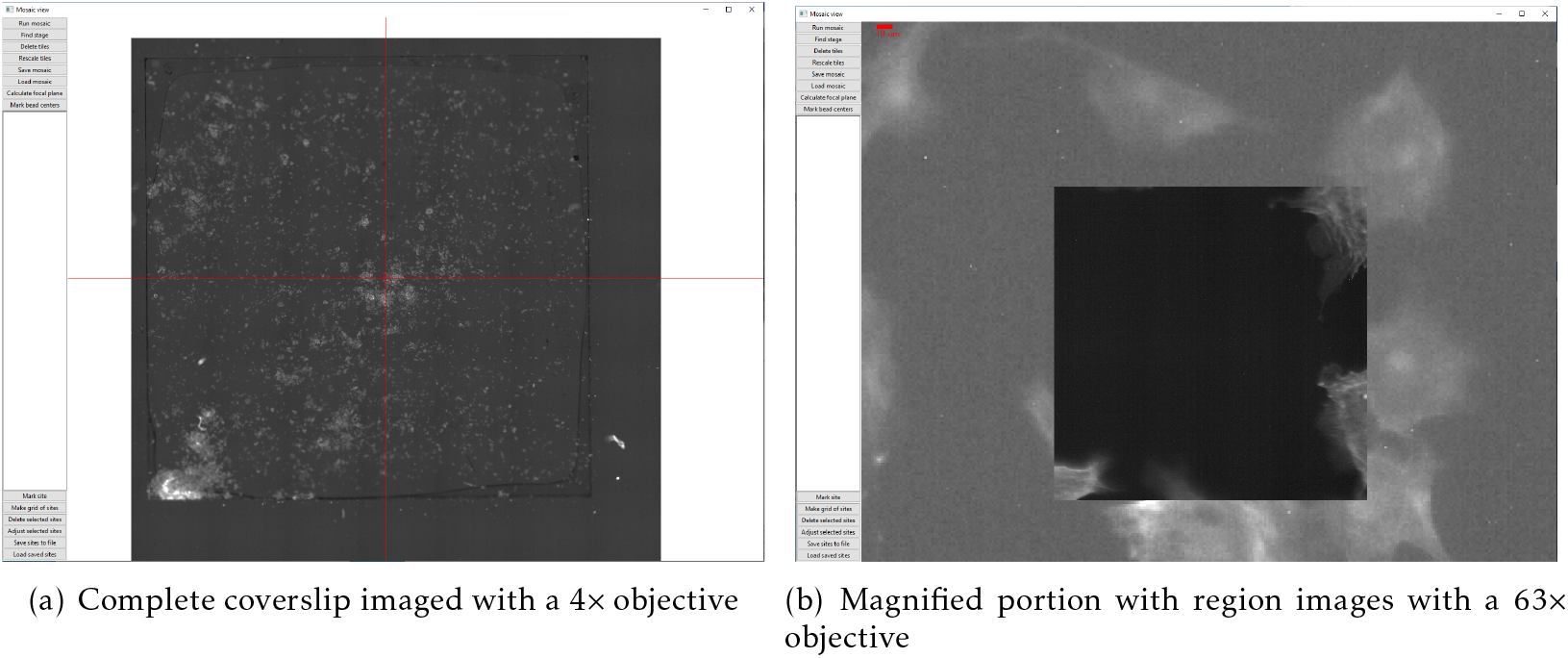
The mosaic tool used to image a standard Thermo Fisher test slide (FluoCells™ Prepared Slide #1, F36924) with: (a) mosaic of a complete coverslip imaged with a 4x objective (pixel size 1.502 μm) consisting of more than 250 megapixels; (b) a magnified portion of the mosaic with a single image from a 63x objective (pixel size 0.0954 μm) painted in on the low resolution map. The higher magnification image has much better signal to noise ratio from better light collection as the objective has a much higher numerical aperture.

To improve the user-centred microscope experience, we divided the use of the microscope into two stages. The user starts an imaging session by rapidly exploring large regions or even the entire slide to create a guide map, while marking regions of interest for later inspection. In the first stage, the touchscreen can be used to build the guide map. The user then establishes the conditions required for the desired images, such as choice of laser lines and their intensities, exposures, number of channels and cameras, Z sectioning range, and points to visit on the guide map. The use of a touchscreen enables rapid setting up of the conditions for the experiment, focusing on the most important features of the microscope hardware without providing full functionality.

In a second phase of acquisition, a more detailed software control window is used with a conventional keyboard to define any more advanced experimental settings, such as numeric values for time lapse interval which are difficult to set from a touch interface. Finally the experiment is run.

In this way, we have created a unique, near-instantaneous control environment that allows the user to start an imaging session by exploring the entire slide quickly and marking regions to visit and image in order to achieve their scientific goals. Our software provides a novel way for the user to plan their entire work flow rapidly and with efficient specimen usage.

### Previously published systems and microscopes in development using Cockpit

To exemplify the flexibility of Cockpit, we present a number of systems that currently run using the software as the user interface. Along with previously published systems, we have a number of microscopes at earlier stages of development where Cockpit provides the user interface.

Cockpit has already been deployed in three systems reported in previous publications. The first is a cryo-super resolution microscope for correlative imaging, CryoSIM [28, 29](Figure 5(a)). The second demonstration was a laser-free spinning disk confocal microscope using adaptive optics (AO) for aberration correction [30] (Figure 5(b)). Additionally, Cockpit was the basis of the control software in the implementation of a novel technique, IsoSense, for improved aberration correction in widefield microscopes and structured illumination [31].

**Figure 5:**
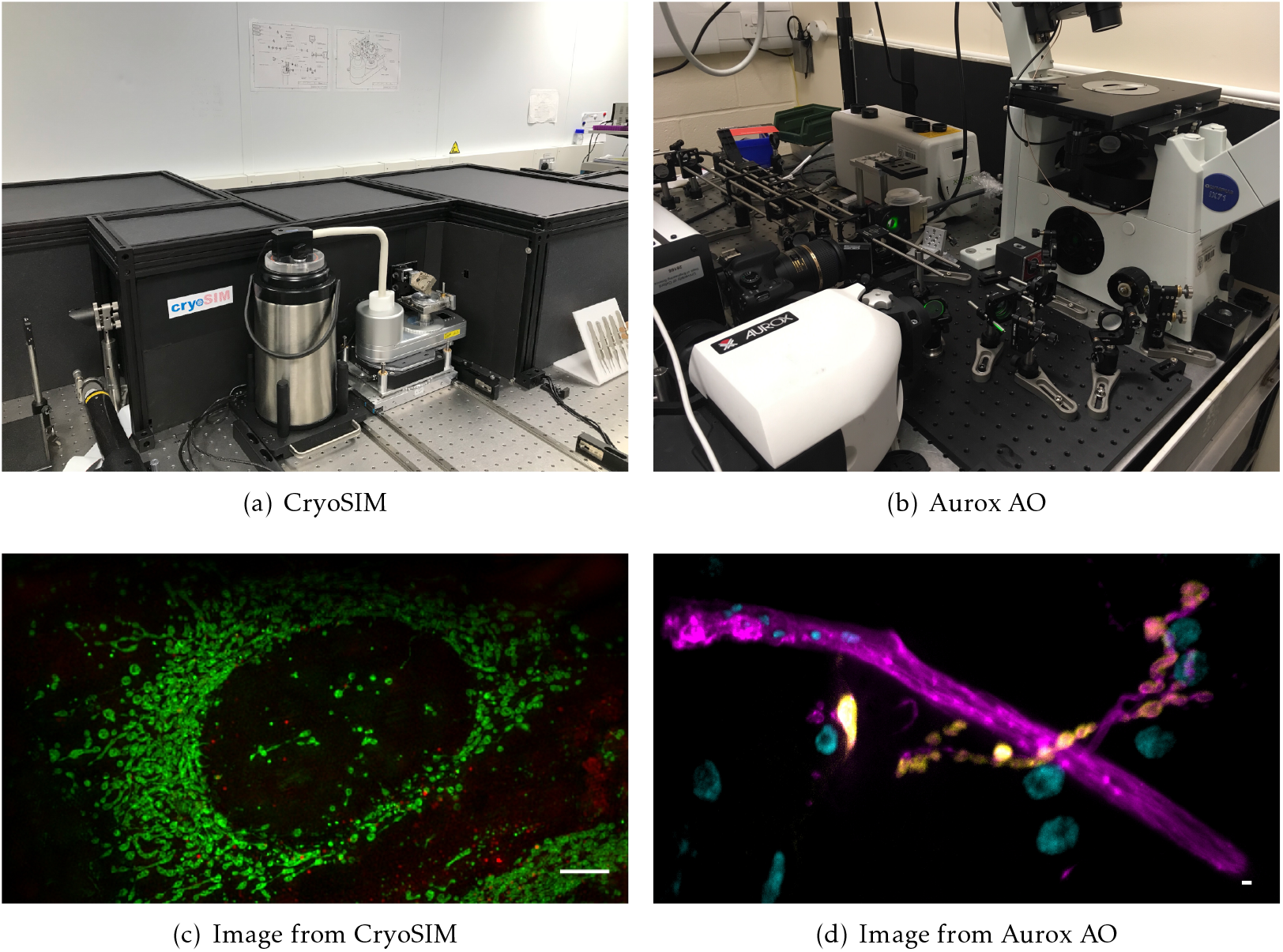
Previously published systems that use Cockpit: (a) the CryoSIM system; (b) the Aurox Clarity AO laser free spinning disk confocal system; (c) image acquired on CryoSIM of cryo preserved HeLa cell labelled with MitoTracker green and LysoTracker red; (d) image acquired on the Aurox Clarity system with AO correction at high optical sectioning. Image is a maximum intensity projection of a 20 μm *z*-stack. Sample was a *Drosophila* neuromuscular junction with DLG in yellow, HRP in magenta, and DAPI in cyan. Scale bars 5 μm.

### A simple widefield microscope

We present a simple, compact, fully automated, portable inverted microscope based around the Zaber MVR system. This provides a bare-bones inverted stand with motorised X, Y, and Z axes, a multi-position motorised filter turret, and LED-based fluorescence illumination. The system has a Ximea camera with a small physical footprint (approximately 30 mm cube) and a Red Pitaya single board computer running as a hardware timing device (Figure 6).

**Figure 6:**
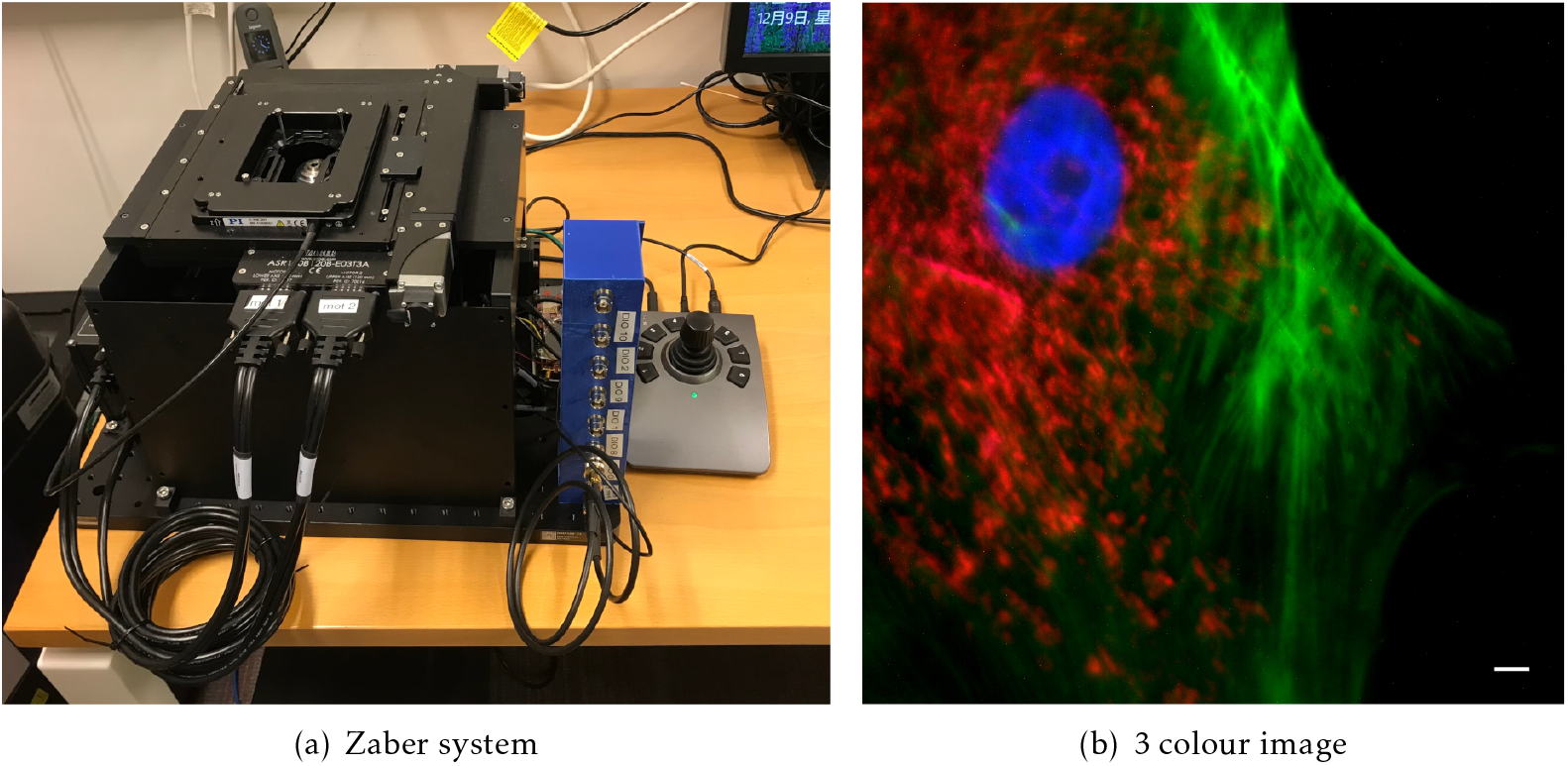
The Zaber microscope. (a) An image of the system, showing its compact size with a 300 Ô 450 mm footprint, (b) a three colour image showing the nucleus stained with DAPI in blue, the actin in green and mitochondria stained with MitoTracker Red CMXRos in Red. Scale bar 5 μm.

This system is able to take fluorescence images in three colours, using the three illumination LEDs and a quad-bandpass dichroic filter set. The system has 4 and 63 air objectives. Although changing the objective involves physically removing the objective and its mount, this is easily achieved due to the spring-loaded kinematic mounting system. The Cockpit mosaic function allows large areas of a sample to be mapped quickly (Figure 4(a)), and then regions of interest can be selected and marked for return later or the mosaic stopped and z-stacks in one or more channels collected. If a live sample is used the system can be used for time lapse imaging.

This system provides a demonstration of the power and portability of Cockpit and it functions as a test bed for new software developments on practical hardware. The system’s small size and portability enable easy transport of the system for demonstrations and collaborative projects in other laboratories.

### A complex structured illumination microscope with adaptive optics

In a further system, we are using Cockpit to maximise the performance of complex upright widefield structured illumination microscope, referred to as DeepSIM (Figure 7(a)). This incorporates not only translation stages, light sources, and cameras, but also a spatial light modulator (SLM) and a deformable mirror (DM) for AO. This set-up is required to perform super-resolution live imaging experiments deep in tissue specimens, such as on the *Drosophila* larval neuro-muscular junction preparation, a powerful model system for understanding synapse biology on a molecular level. In order to achieve this imaging the system has to synchronise the illumination lasers, the SLM, a polarisation rotator, the DM, the Z stage position, and the cameras. The lasers, SLM, DM, and cameras receive digital triggers from the Red Pitaya, while the Z-position and polarisation rotator state are controlled via analogue voltages, all synchronised at the μs level. Cockpit provides a simple user interface allowing selection of different experimental parameters, such as exposure time, laser power, Z step, and stack size (Figure 7(b)). The experiment module creates the relevant signals and timing information, transfers this to the Red Pitaya which then controls the hardware during the experiment.

**Figure 7:**
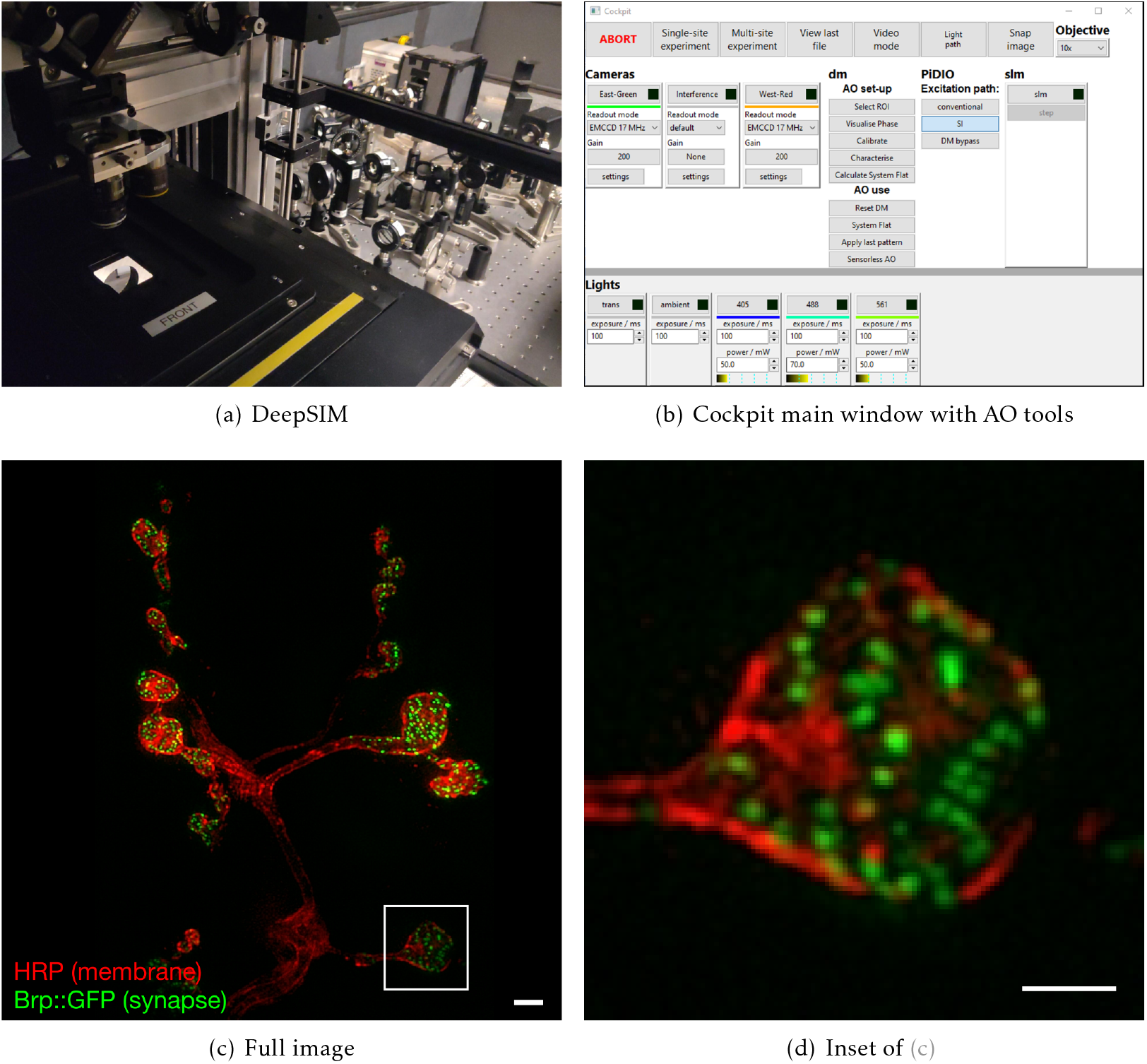
Example data from the DeepSIM system using Cockpit to image deep (*>* 20 μm) in *Drosophila* neuro-muscular junctions. The system uses sensorless AO to correct for sample induced aberrations. (a) system view from the stage; (b) main Cockpit window with the additional controls for the SLM and AO devices; (c) image of individual synapses at the *Drosophila* neuro-muscular junction using AO-SIM imaging, with Cy3 labelled HRP antibody labelling the neuronal membrane in red and Brp::GFP in the synapse in green; (d) zoomed in region of (c). Scale bars are 2 μm.

Cockpit also provides a user-friendly interface to Microscope-AOTools [32], facilitating use of a wavefront sensor for calibration of the DM then adaptively correcting sample induced aberrations via image-based metrics, using so-called sensorless AO. This interface also implements IsoSense[31] to improve aberration detection over a wide range of complex samples.

Although this system is extremely complex with a multitude of devices the user interface is clean and easy to use. The system has sensible defaults, thus minimising the expertise and interaction required to collect experimental data.

## Discussion

We have developed a Python-based GUI for controlling bespoke microscopes which can map the entire slide or dish, allowing a real time exploration of the entire specimen landscape. It also connects to separate hardware timing devices to enable high precision timing of even complex devices and experiments. A key property of our software is that it is focused around the needs of the user and their experimental design and workflow. Finally, using Python means that the microscope control can easily be modified to make use of the extensive machine learning algorithms available in easy to integrate libraries. This means that a microscope system that is controlled by Cockpit can readily be adapted to make decisions using machine learning algorithms during the acquisition process in order to modify the imaging conditions.

### Existing control options

Cockpit has been introduced as an alternative to a number of existing microscope control options that are already available. Most published systems take one of three approaches: 1) utilise individual software packages provided by the device manufactures, 2) use LabVIEW to integrate LabVIEW-based drivers (VIs) provided by the manufacturers into a single custom GUI, or 3) use the open source microscope control package μManager [33]. We will discuss the relative merits of each of these options in turn.

Using individual software packages to control each device is simple and direct. However this approach has several severe drawbacks. Each hardware device needs its own software, which takes up screen space and computer resources. This also removes the ability of different hardware parts to automatically interact with each other, for instance changing a stage’s position once a camera has finished collecting images. Producing Z-stacks, or multi-channel images on such a setup is awkward, slow and open to human errors. These factors can lead to corruption of the results.

LabVIEW is a visual programming tool that allows the construction of “programs” by connecting modules with wires and building graphical interfaces. Many manufacturers provide software to allow their hardware to be used in LabVIEW, through so-called virtual instruments (VIs). Building clean and simple visual framework to control highly complex advanced optical systems in LabVIEW requires advanced expertise in the program’s use. Because such skills are usually beyond the ability of most scientists in an academic setting, many one-off complex bespoke systems, built to demonstrate a new principle, are very hard to use routinely by experimental scientists. Despite their obvious importance, such systems can rarely be reproduced and adopted by others for routine use.

μManager is an open source generalised microscope control interface which works on top of ImageJ, a popular image analysis program. ImageJ is written in Java, and μManager is written in a combination of Java and C++. μManager’s approach is most directly comparable to Cockpit. Many hardware device manufacturers provide μManager compatible libraries and instructions on how to connect and control their hardware. The package comes with mechanisms to run basic experiments, and the ability sequence commands.

Being based on ImageJ, μManager, by design, includes a large application with multiple windows. This also means that the majority of the interface must be written in Java; however the system also needs a C/C++ layer to interface the Java code to C/C++ based system libraries. Cockpit is written in pure Python, relying on the Python-Microscope package for hardware interfacing. Producing a strict separation of the user interface from the hardware control components. This has the additional benefit that connected hardware may be physically located on another computer, increasing scalability and allowing devices requiring incompatible hardware or software to still be seamlessly integrated.

The mosaic window in Cockpit is similar in concept to Micro-Magellan [24], a μManager plugin. Being able to record a large area view, possibly from multiple areas, and have that available for instant navigation. The mosaic functionality dramatically speeds up experiment setup and finding the correct regions of interest, especially on bespoke systems with no eyepieces. Our mosaic interface utilises modern fast GPUs to enable instant access to even very large mosaic maps. Additionally, the touch screen interface allows easy and intuitive access to most of the functionality using a visual grammar that is very familiar to everyone.

The dedicated timing device interface allows fast and repeatable timing for both simple and complex experiments. It is useful in general but absolutely indispensable for experiments like the live cell adaptive optics SIM experiment (Figure 7(c)).

The standardised device interfaces from Python-Microscope mean that replacing one device with another of the same type is simply a matter of changing the address of the device in a configuration file.

## Conclusions

We have developed Cockpit, a new paradigm for user-based software control of complex bespoke microscopes. The software is highly adaptable by a user or engineer with experience in Python. The key advantage of our approach is that a user is presented with a clean and simple interface that hides the complexity of the hardware, so they can focus on their experimental design and obtain data from the instrument with a high precision and reproducible workflow.

## Software availability and development by the community

Cockpit is distributed under the GNU General Public License version 3.0 or later and available from GitHub (https://github.com/MicronOxford/cockpit) a more descriptive website and links at https://micronoxford.com/python-microscope-cockpit.

Cockpit is an open source microscope control software package written in Python. It includes an advanced mosaic mode for experiment setup and sample navigation and the ability to run hardware timed simple and complex experimental protocols. We hope that the community will adopt this package and help us to continue to develop it.

Cockpit is still under active development, both within our labs and at several other laboratories across multiple countries. We are working on adding online image analysis to further enhance the mosaic functionality with machine learning, enabling the system to capture a large area of the sample quickly at low-resolution and then automatically identify features of interest for 3D-multi-channel acquisition.

## Competing interests

Martin Booth declares a significant interest in Aurox Ltd., whose microscopes were used in this work.

## Grant information

This research was funded by Wellcome Strategic Awards and Investigator Award to I.D. [091911/Z/10/Z], [091911/Z/10/A], [105605/Z/14/Z], [107457/Z/15/Z], and [203141/Z/16/Z]; an MRC/EPSRC/BBSRC Next-generation Optical Microscopy [MR/K01577X/1] to I.D.; GIS IBiSA [#2015-28]; and by the CNRS MITI [Défi Imag’In 2015]. D.S. is funded by a BBSRC iCASE grant with Aurox as the industrial partner [grant number BB/M011224/1]. N.H. was supported by funding from the Engineering and Physical Sciences Research Council (EPSRC) and Medical Research Council (MRC) [grant number EP/L016052/1].

## Acknowledgements

We are grateful to the original UCSF software team led by John Sedat, including Melvin Jones, Chris Weisiger and Sebastian Haase for the initial writing of the Python code used to control the OMX and OMX-T microscopes, which was subsequently released as open source software and formed the starting point for this project. We wish to thank many colleagues who have provided helpful suggestions, discussions and testing during development of Cockpit, the Micron Advanced Bioimaging Unit particularly Mantas Žurauskas and Andrew Jefferson, and Maria Harkiolaki from the Diamond Light Source beamline B24.

## Author Contributions

MAP: conceptualisation, software. DMSP: conceptualisation, software, writing — original draft preparation. NH: software. CW: conceptualisation, software. SH: conceptualisation, software. JML: software. RMP: resources. JT: resources DVS: software. TP: software. TSP: software. JWS: conceptualisation. MJB: supervision, conceptualisation. ID: conceptualisation, project administration, writing — original draft preparation, writing — review and editing, supervision, funding acquisition. IMD: conceptualisation, software, writing — original draft preparation, writing — review and editing, supervision, project administration.

## Methods

### CryoSIM imaging

The CryoSIM system has previously been published in detail [28, 29]. It was used to image HeLa cells grown on carbon coated gold EM grids under standard tissue culture conditions. Cells were labelled with MitoTracker Green (Thermo Fisher) at 100 nM and LysoTracker Red DND-99 (Thermo Fisher) at 50 nM after 30 minutes of incubation, to give green mitochondria and red lysosomes. The grids with live cells on them were then blotted and plunge frozen in liquid nitrogen cooled liquid ethane (Leica EM GP2) before being transferred to liquid nitrogen storage. Once frozen, grids were preserved in liquid nitrogen or at cryogenic temperatures on the imaging systems to prevent thawing and detrimental ice crystal formation. Structured Illumination Microscopy (SIM) images were collected with 488 nm and 561 nm laser excitation and emission collected at 525/50 and 605/70 on two Andor iXon EMCCD cameras. Images were reconstructed using SoftWoRx (GE Healthcare) and image quality was assessed with SIMcheck [34].

### Aurox AO imaging

The Aurox Clarity AO system is as previously published, the system utilised an Olympus ix70 microscope with a 60× 1.42NA objective. Illumination was provided by a CoolLED p-300 [30]. This system was used to image *Drosophila melanogaster* neuro-muscular junctions (NMJ). The samples were prepared by following the protocol form Brent et al 2009[35]. 3rd instar *Drosophila melanogaster* larvae (Oregon-R strain) were dissected in HL3 buffer with 0.3*mM* Ca^2+^ to prepare a larval fillet. After this, larvae were fixed with 4% paraformaldehyde and blocked using 1% BSA[35]. Larvae were stained overnight with 1:100 Horseradish Peroxidase (HRP) conjugated to Alexa568 fluorophore to visualise the neurons, and primary mouse antibody against Discs large (DLG) to visualise the postsynaptic density. The next day, the larvae were counter stained with secondary antibody to detect the DLG (1:200 donkey anti-mouse conjugated to Alexa488 fluorophore was used), as well as 1:1000 from 1 mg mL^−1^ stock DAPI to visualise the nuclei. The larvae were then washed and mounted in 65% vectashield.

### Zaber imaging

This system is based around a Zaber MVR motorised inverted microscope with Zeiss optics and 4x, 0.13NA and 63x 0.75 Air objectives. Fluorescence illumination comes via 3 LEDs (385 nm, 473 nm and 568 nm) and a quad bandpass dichroic filter set (Chroma #89402). The system has a small format (approx 30 mm cube) Ximea (model MQ042MG-CM) camera and a Red Pitaya single board computer running as a hardware timing device. Multi channel images were taken by sequentially illuminating with the three different LEDs without changing the quad dichroic filter cube.

### DeepSIM imaging

DeepSIM is a custom made upright structured illumination microscope with AO specifically designed for SIM super resolution imaging deep into live tissue (manuscript in preparation). On the DeepSIM system images were taken using 488 nm and 561 nm laser illumination. The system includes a Structure Light Modulator (SLM, Meadowlark) to provide structured illumination for SIM imaging. The system also includes a deformable mirror (Alpao DM-69) for aberration correction. The system has a 60x 1.1 water dipping objective and samples were images in aqueous buffer. The AO components were calibrated and controlled using Microscope-AOTools [32] to reduce aberrations and produce good SIM imaging at depth in biological samples.

The samples were prepared by following the protocol[35]. 3rd instar *Drosophila melanogaster* larvae (Bruchpilot (Brp)-GFP strain) were dissected in HL3 buffer with 0.3*mM* Ca^2+^ to prepare a so-called larval fillet, and the larval brains were removed. After this, larvae were stained for 15 minutes with 1:50 Horseradish Peroxidase (HRP) conjugated to Cy3 fluorophore to visualise the neurons, washed with HL3 buffer with 0.3*mM* Ca^2+^ and imaged in HL3 buffer with 0*mM* Ca^2+^ to prevent the larvae from moving.

